# Distinct Hypoxia-induced Translational Profiles of Embryonic and Adult-derived Macrophages

**DOI:** 10.1101/2021.03.18.435732

**Authors:** Nicholas S. Wilcox, Timur O. Yarovinsky, Prakruti Pandya, Vinod S. Ramgolam, Albertomaria Moro, Yinyu Wu, Stefania Nicoli, Karen K. Hirschi, Jeffrey R. Bender

**Affiliations:** Department of Internal Medicine, Section of Cardiovascular Medicine, Yale Cardiovascular Research Center, Yale University School of Medicine, New Haven, CT, 06511, United States; Department of Immunobiology, Yale University School of Medicine, New Haven, CT, 06511, United States; Department of Genetics, Yale University School of Medicine, New Haven, CT, 06511, United States

**Author notes:** Authors contributed equally (Nicholas S. Wilcox, Timur O. Yarovinsky). 3400 Spruce Street, Hospital of the University of Pennsylvania, Philadelphia, Pennsylvania, 19104 (Nicholas S. Wilcox).

## Abstract

Tissue homeostasis and repair are orchestrated by resident and newly recruited macrophages that alter their gene expression program in response to changes in tissue microenvironment. Embryonic macrophages, such as fetal liver derived macrophages (FLDM) seed the organs, including heart and lung during embryonic development and persist throughout the adult lifetime, while bone marrow-derived macrophages (BMDM) are recruited following an acute perturbation. Transcriptome analyses of FLDM and BMDM identified differences between them at the level of RNA expression, which correlates imperfectly with protein levels. Post-transcriptional regulation by microRNAs (miRNAs) and RNA-binding proteins determines mRNA stability and translation rate and may override transcriptional cues in response to environmental changes, such as hypoxia. To identify distinct features of FLDM and BMDM response to hypoxia at the level of translation, we employed translating ribosome affinity purification (TRAP) to isolate polysomal RNA. RNA-seq profiling of translated RNA identified distinct hypoxia-induced translational signature of BMDM (Ly6e, vimentin and glycolysis-associated enzymes Pgk1, Tpi1, Aldoa, Ldha) and FLDM (chemokines Ccl7 and Ccl2). By translational profiling of BMDM and FLDM with deletion of the RNA-binding protein HuR, we identified transcripts that were dependent on HuR. These findings highlight the importance of HuR and identify its distinct targets for post-transcriptional regulation of gene expression in embryonic vs. adult-derived macrophages.

## Introduction

Macrophages populate all tissues and exert multiple functions to maintain homeostasis in response to environmental changes (Ginhoux and Jung, 2014). Embryonic macrophages originating from yolk sac or fetal liver populate most organs during early development and constitute the majority of tissue-resident macrophages. Fate mapping studies in mice reveal that more than 95% of cardiac- and lung-resident macrophages originate from fetal liver (Heidt et al., 2014; Lavine et al., 2014). Under homeostatic conditions, tissue resident macrophages undergo self-renewal with little influx from the bone marrow (Epelman et al., 2014). However, organ injury or inflammation results in rapid recruitment of bone marrow-derived monocytes that differentiate into macrophages with overlapping and distinct roles in tissue repair, from those of tissue-resident cells (Dick et al., 2019; Heidt et al., 2014; Lavine et al., 2014; Liao et al., 2018).

Reduced blood flow to organs (i.e., ischemia) decreases oxygen availability in target tissues. The resultant hypoxic conditions are sensed not only by stromal cells, but also by tissue-resident and newly recruited macrophages (Lewis et al., 1999; O’Neill IV et al., 2005). Hypoxia-induced stabilization of hypoxia-inducible factor-1α (HIF-1α), followed by nuclear translocation and association with HIF-1β, leads to transcriptional activation of many genes, including those that enhance ATP production through glycolysis, and those that promote angiogenesis in affected tissues (Rahat, 2011). Posttranscriptional regulation of gene expression at the level of mRNA stability and translation may amplify or override transcriptional regulation and, as such, plays an important role in macrophage responses to hypoxia (Rahat, 2011; Walter et al., 2018).

It is now well established that transcript levels do not always correlate with protein abundance. Multiple RNA-binding proteins, including HuR, PTB, TTP, and others, change mRNA turnover and translation rate (Gorospe et al., 2011). For example, HuR can stabilize macrophage transcripts regulated by HIF-1α, including vascular endothelial growth factor A (Vegf) and matrix metalloproteinase-9 (Mmp9), which modulate angiogenesis and tissue repair (Zhang et al., 2012). Another level of posttranscriptional regulation complexity is added by microRNAs (miRNAs), which cause RNA degradation or translational arrest through binding to mRNA 3’-UTRs (Gorospe et al., 2011). HuR can compete with miRNAs for binding to neighboring 3’-UTR sites, thereby conferring protection (Bhattacharyya et al., 2006; Kundu et al., 2012). The attenuation of miRNA binding to transcripts in close proximity to HuR binding sites, and consequential miRNA targeting of transcripts regulating angiogenesis and macrophage/endothelial interactions, have been demonstrated (Chang et al., 2013). There are global and specific changes in macrophage mRNA translation in response to various stimuli, including hypoxia (Hofmeister et al., 2020; Liu et al., 2014; Rahat, 2011; Shibata et al., 2013). However, prior studies have either focused on bone marrow-derived macrophages (BMDM) or simply did not attempt to address the origin of macrophages. Considering the differences in the differentiation pathways and transcription programs between adult BMDM and embryonically derived macrophages, we hypothesized that BMDM and fetal liver-derived macrophages (FLDM) respond to hypoxia by changing translation of distinct subsets of mRNAs, some in an HuR-dependent manner. These differences might dictate dominant homeostatic and pathologic responses in a given tissue and form the basis for therapeutic targets.

To test our hypothesis, we differentiated macrophages from fetal liver and bone marrow of transgenic mice that express a codon optimized Cre recombinase (iCre) under the control of colony stimulating factor 1 receptor (Csfr1) promoter and, after Cre-mediated recombination, large ribosomal protein L10a fused to EGFP from the Rosa26 locus (Deng et al., 2010; Heiman et al., 2008). This approach enabled us to employ translating ribosome affinity purification (TRAP) followed by RNA-Seq analyses of total vs. translated poly(A) pools in BMDM and FLDM at baseline and after exposure to hypoxia. We found that BMDM, but not FLDM, respond to hypoxia primarily by upregulating translation of mRNAs encoding glycolytic enzymes. Furthermore, BMDM increased translation of multiple functionally diverse immune regulators, whereas FLDM increased translation of CC chemokines Ccl7 and Ccl2. Finally, by using conditional deletion of HuR in macrophages, we identified transcripts that were differentially regulated by this RNA-binding protein in the context of hypoxia.

## Results

### BMDM and FLDM for translational profiling

To compare mRNA translation in macrophages of embryonic vs. adult origin, we expanded macrophages from fetal livers or adult bone marrow of Csf1r^iCre^Rosa26^EGFP:L10a^ mice in the presence of conditioned medium from L929 fibroblasts containing M-CSF for 6-7 days. More than 90% of the cells in culture were GFP+, indicating expression of EGFP-L10a, and more than 90% of GFP+ cells were positive for CD11b and F4/80, a marker for differentiated macrophages (Figure 1A). BMDM and FLDM cultures were similar morphologically with EGFP-L10a present in both the nucleus and cytoplasm (Figure 1B).

**Figure 1.**
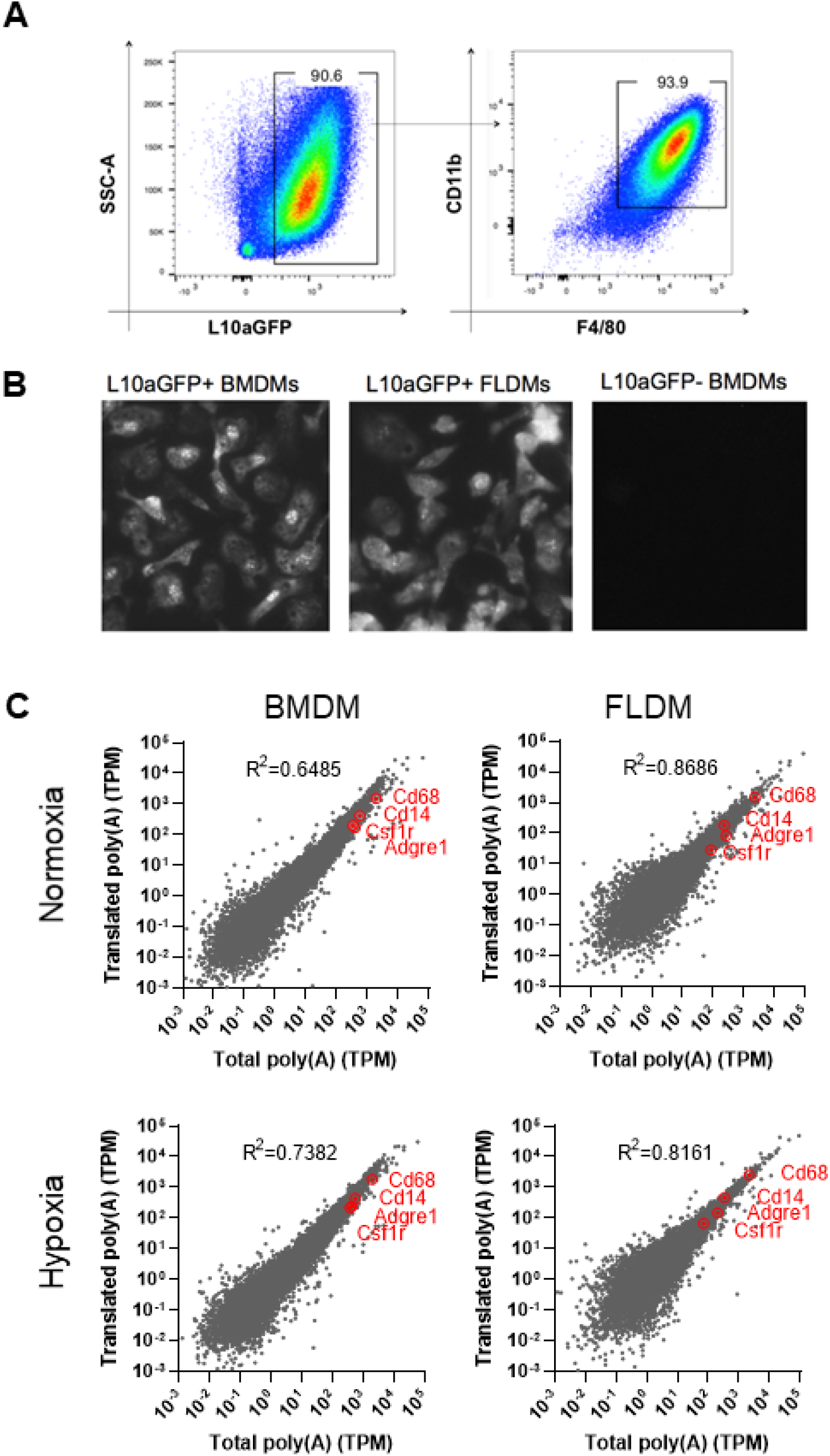
Characterization of BMDM and FLDM phenotype, expression of EGFP-L10 and total vs. translated poly(A) RNA pools at baseline (normoxia) and after exposure to hypoxia. (A) Representative FACS plots reveal that more than 90% of L10aGFP+ cells in FLDM cultures are F4/80+CD11b+ by day 6 of culture in the presence of M-CSF. Similar pattern of staining was routinely observed in BMDM cultures. (B) Representative fluorescence microscopy images display EGFP-L10a expression in the differentiated BMDMs and FLDMs from Csf1r^iCre^L10aGFP+ mice compared to L10aGFP^-^ control. (C) Normalized results of RNA-Seq (TPM) of total vs. translated poly(A) pools for both macrophage types under normoxic and hypoxia conditions. The data are mean of 3 (BMDM) or 2 (FLDM) independently processed samples. Red encircled dots indicate the abundance and translation of mRNA encoding macrophage markers Adgre1 (F4/80), Csf1r (c-fms, or colony stimulating factor 1 receptor), CD14 and CD68.

We exposed differentiated BMDM and FLDM to hypoxia (1% O_2_, 5% CO_2_, balanced with nitrogen) for 18 h and applied RNA-Seq analyses to the total vs. translated poly(A) RNA pools isolated with translated ribosome affinity purification (TRAP). The results were expressed as Transcripts Per kilobase Million (TPM) to determine abundance and translation for each mapped transcript. The mRNAs encoding macrophage-specific markers Adgre1 (F4/80), Csf1r, CD14 and CD68 were highly abundant in total and translated RNA pools for both macrophage types under normoxic and hypoxic conditions (Figure 1C). At whole transcriptome level, strong correlation was evident between abundance and translation of most transcripts in BMDM and FLDM under normoxia and hypoxia conditions (p<0.0001) (Figure 1C). However, a number of transcripts were either under-represented or over-represented in the translated poly(A) relative to total poly(A) pools with apparently distinct patterns between BMDM and FLDM (Figure 1C).

### Hypoxia-induced changes in the total poly(A) RNA transcriptome

RNA-Seq analyses of the total poly(A) RNA showed that exposure to hypoxia resulted in significant upregulation or downregulation of 22 protein-coding transcripts in BMDM and 24 protein-coding transcripts in FLDM, with 8 of those transcripts changing significantly in both cell types (Figure 2A). Within the shared subset, 3 mitochondrially-encoded protein coding transcripts (mt-Nd1, mt-Nd2, and mt-Cytb) were downregulated, whereas the nucleus-encoded transcripts were either upregulated in both cell types (Pf4 and Tmsb10) or changed in the opposite directions (Eef1a1, Prdx1, Ccl7) (Figure 2B). Specific for BMDM response to hypoxia were significant upregulation of the transcripts encoding enzymes involved in glycolytic process (Pgk1, Tpi1, Aldoa, Pkm, and Ldha) and transcripts related to regulation of innate immunity (C5ar1, Mif, Vim, S100a4, Saa3), as well as changes in the transcripts related to lipid metabolism (Apoe, Lpl) and down-regulation of a mitochondrially-encoded transcript mt-Nd5. FLDM responded to hypoxia by upregulation of transcripts encoding CC-chemokines (Ccl2, Ccl7, Ccl4), lysozyme 2 (Lyz2), Fe and Zn transport proteins (Crip1, Flt1), and actin-binding proteins (Pfn1, Tmsb4x). Remarkably, while Ccl2 and Ccl7 topped the list of most upregulated transcripts in FLDM, Ccl2 did not change and Ccl7 was the most downregulated transcript in BMDM (Figure 2B). Specific for FLDM were also changes in the abundance of several transcripts related to protein synthesis (encoding ribosomal proteins Rps27l, Rpl32, Rpl18a) and folding (Ppia), while transcripts for actin (Actb), fatty acid-binding protein (Fabp5) and a subset of mitochondrially-encoded transcripts (mt-Co1, mt-Nd-6 and mt-Nd-4) were downregulated. These results indicate that hypoxia responses in BMDM and FLDM can be characterized by a common pattern in downregulation of mitochondrially-encoded transcripts, but differential changes in the abundance of nucleus-encoded transcripts associated with innate immunity and cell metabolism, including glycolysis and protein synthesis.

**Figure 2.**
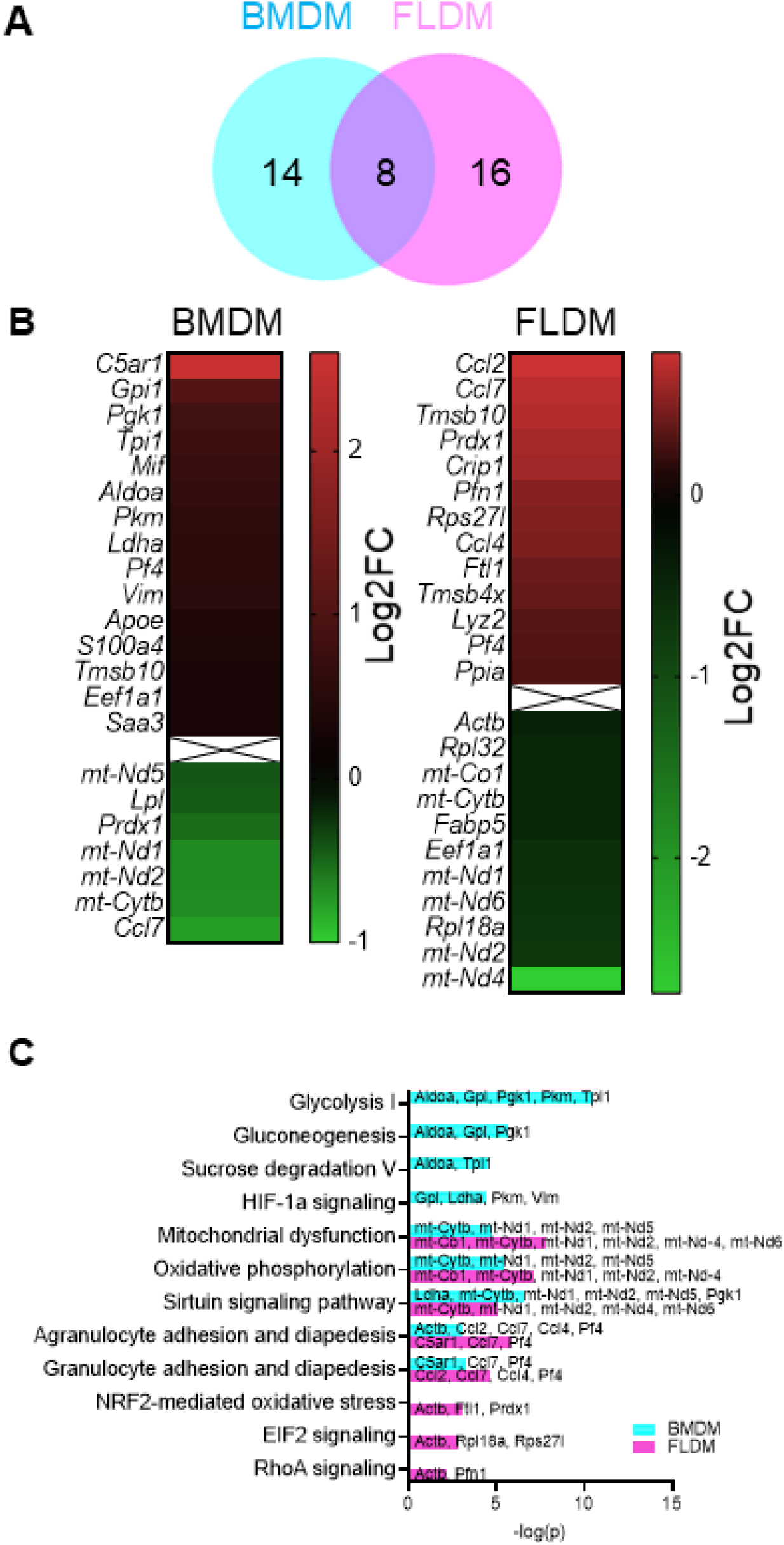
Analyses of changes in the total mRNA abundance in BMDM and FLDM in response to hypoxia. (A) Venn diagram showing the number of specific and shared mRNAs abundance of which significantly changed in hypoxia. (B) Heat map of the mRNAs that significantly increased or decreased in abundance in each macrophage subset. The data are mean Log2 fold change. (C) Canonical pathways identified by analyses of the hypoxia-induced changes in total mRNA in BMDM and FLDM.

Canonical pathway analyses of hypoxia-induced changes in total mRNA expression revealed pathways that were affected in BMDM or FLDM only as well as a subset of shared pathways (Figure 2C). While activation of glycolysis I (z-score=2.236) and HIF-1α signaling (z-score=2) were specific for BMDM, oxidative phosphorylation showed marked inactivation in both types of macrophages (z-score=−2 in BMDM and −2.236 in FLDM). Mitochondrial dysfunction and sirtuin signaling were similarly disturbed in both types of macrophages, mainly due to downregulation of mitochondrially encoded mRNAs, although z-scores for these pathways did not show significant directional change. Hypoxia-induced changes in C5a1 and chemokine mRNA expression were linked to disturbance of granulocyte and agranulocyte adhesion and diapedesis pathways in both macrophage types, albeit this was due to different transcript subsets and did not show significant directional change. Gluconeogenesis and sucrose degradation pathways were disturbed in BMDM only, while NRF2-mediated oxidative stress, EIF2 signaling and RhoA signaling pathways were disturbed in FLDM only, although without directional change.

### Hypoxia-induced changes in translated poly(A) RNA

RNA-Seq analyses of the translated poly(A) RNA samples showed that exposure to hypoxia resulted in significant modulation of translation of 27 transcripts in BMDM and 9 transcripts in FLDM, with 3 transcripts overlapping between the two cell types (Figure 3A). Within the shared subset, translation of the transcripts encoding transmembrane immune signaling adapter (Tyrobp) and lysozyme 2 (Lyz2) were significantly upregulated, whereas translation of β2-microglobulin (B2m) transcript was upregulated in BMDM and downregulated in FLDM. Increased translation of transcripts encoding enzymes involved in glycolysis (Pgk1, Tpi1, Ldha, and Aldoa) and some of the previously noted transcripts related to regulation of innate immunity (Vim, Mif, S100a) continued to be specific for BMDM (Figure 3B).Moreover, translation of several additional transcripts related to immune regulation (Ly6e, Ifitm2, Pf4, Fcgr3, Wfdc17), CC-chemokines (Ccl6 and Ccl5), carbohydrate binding (Lgals1), actin binding (Tmsb10), and posttranslational protein modifications (Sh3bgrl3, Itm2b, Ctsss) increased specifically in BMDM. In addition, translation of two transcripts encoding large ribosomal proteins changed in opposite directions in BMDM: increased for Rpl19 and decreased for Rplp2. Translation of three other transcripts (Cd63, Prdx1, Calr) was also decreased in BMDM. Hypoxia-induced changes specific for FLDM comprised of increased translation of CC-chemokines Ccl2 and Ccl5 and decreased translation of a component of IgE receptor (Fcerg1), galactose-specific lectin binding IgE (Lgals3), ferritin heavy chain (Fth1), and cystatin-B (Cstb) (Figure 3B).

**Figure 3.**
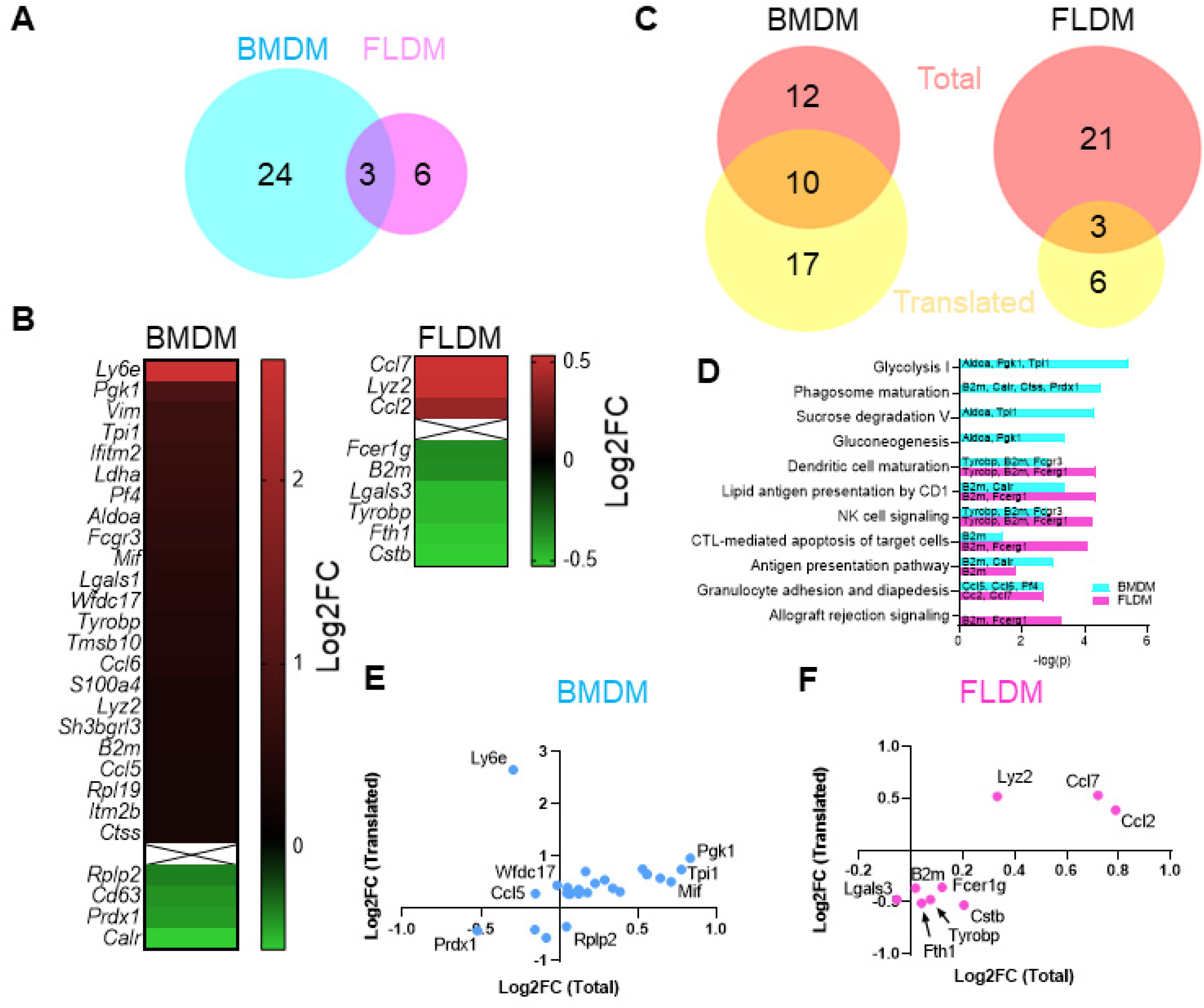
Analyses of the changes in translated mRNA in BMDM and FLDM in response to hypoxia. (A) Venn diagram showing the number of specific and shared mRNAs translation that significantly changed in hypoxia. (B) Heat map of the mRNAs that displayed significantly increased or decreased translation in each macrophage subset. The data are mean Log2 fold change. (C) Venn diagrams showing the number of mRNAs that changed in abundance only, translation only, or in both readouts, for each macrophage subset. (D) Canonical pathways identified by analyses of the hypoxia-induced changes in translated mRNAs in BMDM and FLDM. (E) and (F). Change in abundance (x-axis) and translation (y-axis) are plotted for mRNAs that showed significant changes in translation BMDM (E) and FLDM (F).

Canonical pathway analyses of hypoxia-induced changes in translated mRNAs associated responses to hypoxia with pathways for glycolysis I, phagosome maturation, sucrose degradation and gluconeogenesis in BMDM and allograft rejection signaling in FLDM (Figure 3D). Several pathways (dendritic cell maturation, lipid antigen presentation by CD1, NK-signaling, CTL-mediated apoptosis of target cells, antigen presentation, and granulocyte adhesion and diapedesis) were shared by BMDM and FLDM, although they were represented by few transcripts that were either regulated in different directions (translation of Tyrobp and B2m increased in BMDM and decreased in FLDM) or were distinct (Fcgr3, Calr, Ccl5, Ccl6, PF4 in BMDM vs. Fcerg1, Ccl2, Ccl7 in FLDM).

Next, we determined whether hypoxia-induced changes in mRNA translation could be due to the corresponding changes in the transcript abundance or regulation at the level of translation. In BMDM, significant changes in translation of 10 out of 27 transcripts were accompanied by significant changes in their abundance, whereas changes in translation for the remaining 17 transcripts occurred without significant changes in their abundance (Figure 3C). In FLDM, changes in translation of 3 out of 9 transcripts were accompanied by significant changes in their abundance (Figure 3C). By plotting Log2FC for hypoxia-induced changes in total vs. translated poly(A) RNA, we analyzed the direction of the changes for each transcript (Figures 3E). In BMDM, it is evident that the direction of translation changes is the same as the direction of the changes in the abundance for most transcripts, even though they were not always statistically significant. However, translation of Ly6e and Ccl5 significantly increased while their abundance in the total poly(A) pool showed a trend towards decrease, albeit without statistical significance, whereas translation of Wfdc17 increased and translation of Rplp2 decreased with minimal difference in their abundance. In FLDM, increased translation of Ccl7, Ccl2, and Lyz2 was accompanied by significant increases in the corresponding transcript abundance, whereas decreased translation of Lgals3 was accompanied by a trend towards a decrease in its abundance (Figure 3F). However, significant downregulation of translation of other transcripts occurred despite a trend toward their increased abundance (Fcerg1, Cstb, Tyrobp, Fth1, B2m). Thus, while changes in the translation of many mRNAs may be explained by their changes in their abundance, increased translation of Ly6e, Ccl5, and Wfdc17 in BMDM and decreased translation of Fcerg1, Cstb, Fth1, and B2m in FLDM apparently occur independently of the changes in the corresponding transcripts.

### Effects of HuR deletion on hypoxia-induced changes in mRNA abundance and translation in BMDM and FLDM

To determine whether HuR regulates translation responses to hypoxia in BMDM and FLDM, we analyzed hypoxia mRNA profiles in BMDM and FLDM isolated from Csf1r^iCre^Rosa26^EGFP-L10a^ mice crossed to HuR^fl/fl^ mice (Zhang et al., 2012). Efficiency of HuR deletion in cultured macrophages exceeded 94% and hypoxia had minimal effects on HuR expression in WT cells (Figure 4A). Deletion of HuR had a profound impact on the hypoxia-induced changes in mRNA abundance and translation. Of the 22 transcripts that were either upregulated or downregulated by hypoxia in wild-type BMDM, only 8 transcripts (Tpi1, Pkm, Ldha, Saa3, Lpl, mt-Nd1, mt-Nd2, and mt-Cytb) were significantly regulated by hypoxia in HuR-KO BMDM (Figure 4B). While hypoxia resulted in significant changes in translation of 27 transcripts in wild-type BMDM, translation of only 6 transcripts (Ly6e, Mif, Lgals1, Tmsb10, Ccl5, and Prdx1) in HuR-deleted BMDM was altered by exposure to hypoxia. Of the 24 transcripts that were either upregulated or downregulated by hypoxia in wild-type FLDM, only 7 transcripts (Ftl1, Lyz2, mt-Cyb, Eef1a1, mt-Nd6, Rpl18a, and mt-Nd2) were significantly regulated by hypoxia in HuR-deleted FLDM. Remarkably, none of the 9 transcripts that were differentially translated in response to hypoxia in FLDM showed statistically significant changes in translation in HuR-deleted FLDM. These results suggest that hypoxia-induced changes in mRNA abundance and translation are dependent to a large extent on HuR.

**Figure 4.**
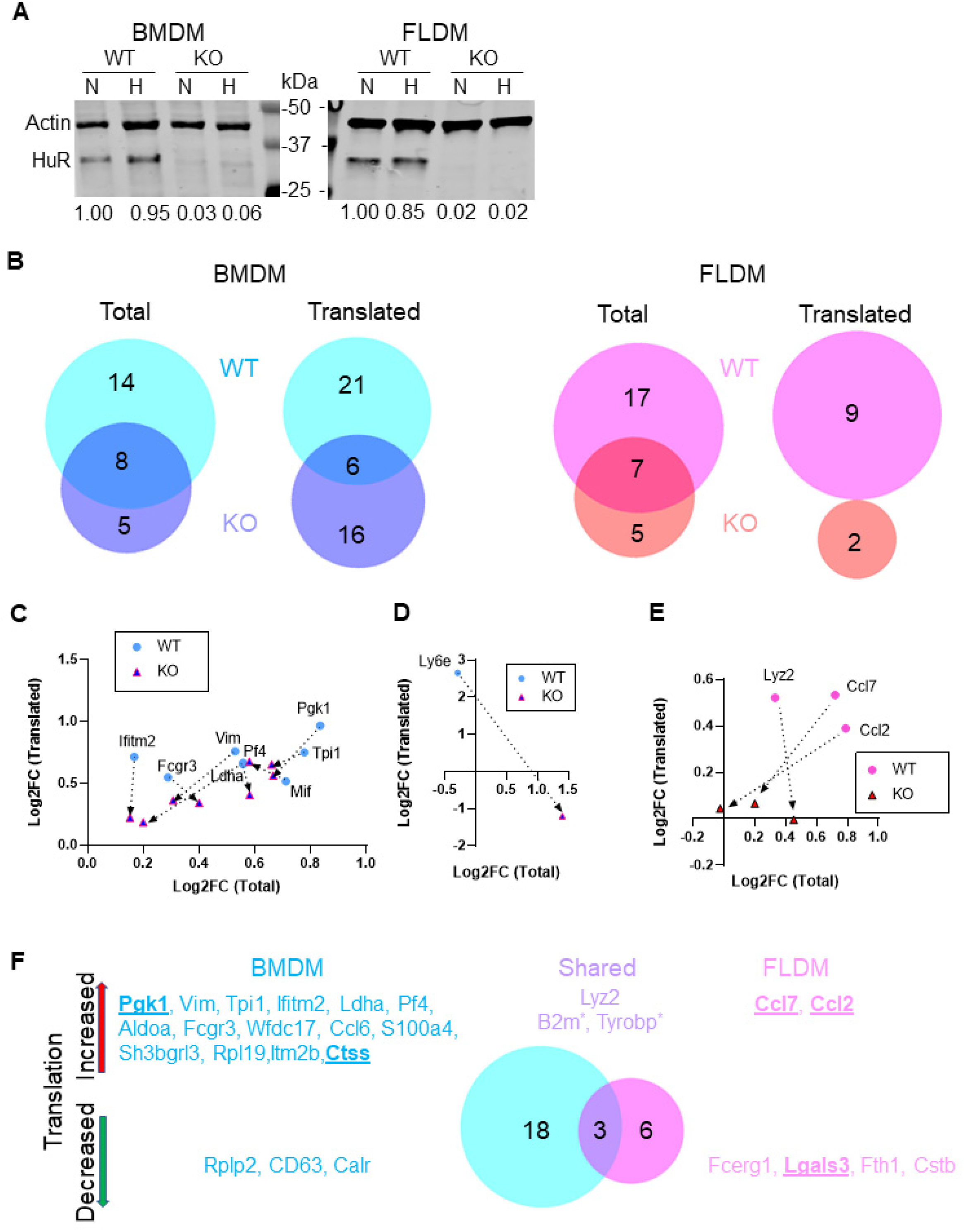
Effects of HuR deletion in macrophages on abundance and translation of mRNA transcripts regulated by response to hypoxia. (A) Representative immunoblot analyses of lysates from BMDM and FLDM isolated fom HuR^+/+^Csf1r^iCre^L10aGFP^+^ (WT) and HuR^fl/fl^Csf1r^iCre^L10aGFP^+^ (KO) mice and exposed to hypoxia. (B). Venn diagrams showing the number of mRNAs that changed in abundance or translation in WT, KO, or both macrophages. (C) and (D). Effects of HuR deletion on select transcripts that showed significant increase in translation in BMDM. (E) Effects of HuR deletion on transcripts that showed significant increase in translation in FLDM. (F) BMDM- or FLDM-specific or shared mRNAs showing hypoxia-induced change in translation in HuR-dependent mode. Previously described HuR targets are underlined in bold font. Asterisk (*) indicate increased translation in BMDM, but decreased translation in FLDM.

Next, we analyzed the direction of the HuR-deletion effect on responses to hypoxia by focusing on transcripts that showed increased translation in wild-type macrophages. In BMDM, hypoxia-driven increases in both abundance and translation of Pgk1, Tpi1, Pf4, and Vim were blunted by HuR deletion (Figure 4C). On the other hand, deletion of HuR resulted in slightly higher increase in Mif translation and lower increase in its abundance. Furthermore, HuR deletion slightly blunted increases in translation of Fcgr3, Ldha, and Ifitm2, but had an opposite effect on hypoxia-induced changes in Fcgr3 and Ldha mRNA abundance. Most striking was the effect of HuR deletion on hypoxia-induced changes in Ly6e expression: although wild-type BMDM responded to hypoxia by increasing Ly6e translation with a trend towards decreased abundance, HuR-deleted macrophages responded to hypoxia by reduction of Ly6e translation and dramatic increase in abundance (Figure 4D). In FLDM, HuR deletion blunted hypoxia-induced increases in Ccl7 and Ccl2 translation and abundance, but blunted translation of Lyz2 mRNA with minimal impact on its abundance (Figure 4E).

In aggregate, RNA-Seq translational profiling of wild-type and HuR-deleted BMDM and FLDM identified distinct mRNAs that are regulated by HuR in response to hypoxia (Figure 4F). Within the shared subset, translation of Lyz2 increased in both cell populations, whereas translation of B2m and Tyrobp increased in BMDM, but decreased in FLDM. BMDM-specific HuR-regulated subset of transcripts encode enzymes involved in glycolysis, ribosomal proteins and regulators of immune responses. All FLDM-specific HuR-regulated transcripts identified in this study encode proteins with described immune functions. Five transcripts identified in this study have been previously shown to be targets of HuR regulation in other cell types: Pgk1, Ctss, Ccl2, Ccl7, and Lgals3 (Burkhart et al., 2013; Fan et al., 2011; Lin et al., 2017; Stellos et al., 2016). Other transcripts may represent novel targets of regulation by HuR.

## Discussion

We characterized total and translated poly(A) RNA in differentiated macrophages of embryonic (FLDM) and adult (BMDM) origin at baseline and after exposure to hypoxia at the whole transcriptome level. We expected to observe both similarities and distinctions between BMDM and FLDM in translated mRNAs. Indeed, the mRNAs encoding macrophage-specific markers F4/80/Adgre1, Csfr1, CD14, and CD68 were expressed and translated at high levels and did not change in response to hypoxia in BMDM and FLDM. Translation of the myeloid lysozyme M (Lyz2) increased after hypoxia in both macrophage types. BMDM and FLDM responded similarly to hypoxia by decreasing abundance of mitochondrially encoded mRNAs in the total poly(A) RNA. However, only augmented expression of transcripts encoding glycolytic enzymes, both at the level of total poly(A) RNA and of translation was observed only in BMDM. Furthermore, BMDM and FLDM showed striking differences in the induction of chemokines and several other transcripts encoding immune regulators. While BMDM increased translation of Ly6e, Ifitm2, Pf4, Fcgr3, Mif, Tyrobp, Ccl6, and Ccl5, and decreased translation of Calr, FLDM increased translation of Ccl2 and Ccl7 and decreased translation of Fcer1g, Lgals3, Tyrobp and Fth1. Overall, these findings support our hypothesis that embryonic and adult macrophages respond to hypoxia by changing translation of distinct mRNA subsets.

Application of the L10a-GFP TRAP assay enabled comparative analyses of actively translated mRNAs. Given abundant posttranscriptional regulatory influences on mRNA stability and translation, we expected differences between hypoxia-induced total and actively translated mRNA profiles. Indeed, changes in mRNA translation and abundance were opposite for some transcripts (Figure 3). Increased or decreased translation in response to hypoxia could be explained by changes in mRNA abundance for approximately a third of transcripts (10/27 in BMDM and 3/9 in FLDM). Some of the differentially translated transcripts showed similar trends in the changes of the total poly(A) abundance but did not reach statistical significance. However, some the of the differentially translated transcripts showed opposite trends in their abundance. Most notably, translation of Ly6e mRNA in BMDM significantly increased despite the trend towards its decreased abundance. At the same time, translation of Cstb and Fcerg1 in FLDM decreased despite the trend towards decreased abundance. These observations suggested that separate mechanisms regulating mRNA abundance and translation may be engaged in response to hypoxia. Further studies will be necessary to determine the exact nature of the upstream regulators and their point of action, such as regulation of transcription or mRNA turnover vs. translational control.

In this study we examined how the RNA-binding protein HuR regulates mRNA abundance and translation in macrophage responses to hypoxia (Figure 4). Using the same Cre driver as for the L10a-EGFP expression, we deleted HuR in differentiated macrophages and carried out the L10a-GFP TRAP assays followed by RNA-Seq. By comparing the hypoxia-induced changes in total and translated poly(A) in wild-type vs. HuR-deleted macrophages, we identified transcripts that required HuR for differential translation (Figure 4F). While several transcripts, such as Pgk1, Ctss, Ccl2, Ccl7, and Lgals3, were previously described as direct targets of HuR in cell types other than macrophages (Burkhart et al., 2013; Fan et al., 2011; Lin et al., 2017; Stellos et al., 2016), this study is the first to describe how they are regulated by HuR in embryonic vs. adult derived macrophages in response to hypoxia. Moreover, we identified 22 additional potential targets of HuR-dependent post-transcriptional regulation. In the future, we will investigate whether these mRNAs are direct targets of HuR or regulated by HuR indirectly through regulation of HuR-dependent genes.

Detection of mitochondrially-encoded transcripts in total poly(A) fractions (Figure 2) was not surprising, given that mammalian mitochondrial mRNAs are post-transcriptionally polyadenylated with approximately 55 A residues by mitochondrial poly(A) polymerase (Fernández-Silva et al., 2003). The translated poly(A) pools also contained mitochondrially encoded mRNAs, albeit at much lower amounts that did not change significantly in response to hypoxia. This may have been the result of contamination of the TRAP reactions by mitochondrial mRNAs, due to their very high abundance in macrophages. The L10a protein, used for the EGFP-L10a TRAP to isolate nucleus-encoded translated poly(A) RNA, is a component of the 60S subunit, one of the subunits of the 80S ribosome (Heiman et al., 2008). Mitochondrial mRNAs are translated by mitochondrial 55S ribosome without involvement of 80S ribosome (Fernández-Silva et al., 2003). Hypoxia-induced downregulation of multiple mitochondrially-encoded transcripts within the total poly(A) RNA pool in BMDM and FLDM most likely represents mitochondrial dysfunction, as a universal response of the mitochondrial transcriptome to hypoxic conditions (Jian et al., 2010).

While mitochondrial dysfunction in response to hypoxia was shared by BMDM and FLDM, hypoxia-induced activation of glycolysis pathways, as evidenced by increased abundance and translation of several mRNAs encoding glycolytic enzymes, was clearly preferential for BMDM and indicates metabolic reprogramming. It remains to be determined whether translation of glycolysis-related mRNAs in FLDM at baseline is sufficient to provide for increased glycolytic demands under hypoxic conditions. It is possible that activation of the HIF-1α pathway by hypoxia in BMDM leads to transcriptional activation of *Pgk1, Tpi1, Aldoa, and Ldha* genes, with increased abundance of the corresponding mRNAs leading to increase in their translation. We cannot rule out a possibility that the apparent dependence of increased Pgk1, Tpi1, Aldoa, and Ldha on HuR expression is due, at least in part, to indirect effects of HuR on gene transcription, such as previously described stabilization of Hif1a mRNA by HuR. However, decreased fold change in Ldha translation in HuR-KO BMDM, despite a trend towards increased Ldha mRNA abundance (Figure 4C), suggests regulation at the level of mRNA translation that is specific for BMDM.

Hypoxia-induced changes in BMDM and FLDM mRNA translation included a number of mRNAs encoding immune regulators, although the repertoire of those mRNAs was different between the macrophage types. Increased translation of RNAs encoding CC-chemokines Ccl7 and Ccl2 in FLDM is consistent with the previously described induction of alveolar macrophage Ccl2 in response to hypoxia *in vivo* (Chao et al., 2011), since alveolar macrophages are largely FLDM-derived (Guilliams et al., 2013; Schneider et al., 2014). These findings suggest that macrophages of embryonic origin may orchestrate leukocyte recruitment and inflammation in the context of hypoxia. Ultimately, dissecting competitive influences between HuR and repressive miRNAs, at 3’-UTRs of RNAs encoding both injury-promoting and repair-promoting gene products, will allow targeted approaches that favor homeostasis or at least minimize injury.

## Materials and Methods

### Animals and macrophage differentiation

We purchased Csf1r^iCre^ (FVB-Tg(Csf1r-icre)1Jwp/J, stock # 021024) and Rosa26^EGFP:L10a^ (B6;129S4-Gt(ROSA)26Sortm9(EGFP/Rpl10a)Amc/J, stock # 024750) from the Jackson Laboratory and backcrossed them to C57BL/6J genetic background to a total of 6 generations prior to establishing the cross of mice homozygous for both Csf1r^iCre^ and Rosa26^EGFP:L10a^ transgenes. To delete HuR in differentiated macrophages, we crossed Csf1r^iCre^Rosa26^EGFP:L10a^ mice to HuR^fl/fl^ mice that were previously generated and maintained on C57BL/6J genetic background in our laboratory (Zhang et al., 2012). We used the resulting HuR^fl/fl^Csf1r^iCre^Rosa26^EGFP:L10a^ mice (8-12 week-old) to isolate HuR-KO BMDM or to establish timed breeding for isolation of embryonic HuR-KO FLDM. HuR^+/fl^Csf1r^iCre^Rosa26^EGFP:L10a^ mice were used for control BMDM and HuR^+/+^Csf1r^iCre^Rosa26^EGFP:L10a^ mice were used for control FLDM. All animal procedures were approved by Yale University IACUC.

We used previously established protocol for in vitro differentiation of BMDM from femur and tibia bones from 8-12 week-old mice that were euthanized by cervical dislocation following anesthesia with isoflurane (Davies and Gordon, 2005). We differentiated FLDM from single cell suspension obtained from E14.5 embryonic livers. We confirmed the date of conception (D0) by identification of vaginal mucus plug in dam and tracked progression of successful pregnancy by tracking changes in body weights. Dams were anesthetized by isoflurane and euthanized by cervical dislocation. Embryonic age was confirmed by counting somite pairs in embryos after dissection. Fetal livers were dissected and pooled from one litter into 10 ml of PBS supplemented with 10% FBS, mechanically dissociated by pipetting up and down vigorously, before being filtered through a 40-μM cell strainer into a 50-mL conical tube and washed by centrifugation (1500 rpm at 4°C for 5 minutes). To remover the contaminating red blood cells, we resuspended the pellets in 10 mL of ACK lysis buffer (0.15 mM NH4Cl, 1 mM KHCO3, 0.1 mM EDTA) and incubated for 5 minutes prior to washing in 20 mL of FBS-supplemented PBS. Differentiation of BMDM and FLDM was started from 3.5*10^6^ cells/plate in 100-mm non-tissue culture treated plastic Petri dishes in 10 ml of RPMI-1640 culture media supplemented with 10% FBS, 2mM L-glutamine, 100 U/ml penicillin/streptomycin, 10 mM HEPES, 50 μM 2-mercaptoethanol and 33% L-cell conditioned medium and proceeded for 6 days under identical conditions. Fresh culture medium was added at day 3. Non-adherent cells were removed with change of medium at day 6. Macrophage differentiation and L10a-EGFP expression were confirmed by observation of adherent cells and EGFP fluorescence using inverted microscope (Leica DM IRB) equipped with DC350FX camera (Leica) and ImagePro software as well as by expression of macrophage-specific marker (F4/80) using flow cytometry.

### Hypoxia exposure

On day 7 of culture, cells were placed in a hypoxic modular incubator chamber (Billups-Rothenberg). Petri dishes containing cultured cells were placed in the chamber. A ring clamp was used create an air-tight seal along the O-ring of the chamber. The tubing of the outlets was unclamped and connected to a tank containing a mixture of 1% O_2_ and 5% CO_2_ balanced with nitrogen (Airgas East). The chamber was gassed for 10-minutes before the tubing was disconnected from the tank and outlets quickly clamped. The hypoxia chamber was subsequently placed in a 37°C incubator. Normoxic controls were concurrently incubated at 37°C. The duration of hypoxia was 18 h and the chamber was re-gassed at 4-hour intervals.

### Translating ribosome affinity purification (TRAP) and RNA isolation

We obtained the total and translated RNA pools using previously published protocol with minor modifications (Heiman et al., 2014). At the conclusion of the 18-hour incubation under normoxia or hypoxia conditions, cells were treated with 100 μg/ml cycloheximide for 30 min, washed with ice-cold PBS containing 100 μg/ml cycloheximide, and lysed in ice-cold lysis buffer (150 mM KCl, 10 mM MgCl2, 20 mM HEPES, 1% NP-40, 0.5 mM DTT, 100 μg/ml cycloheximide) supplemented with Complete Protease and Phosphatase inhibitors (Roche), and 2000 U/ml RNaseOUT™ (ThermoFisher). Scraped lysates were precleared by centrifugation at 2000 g for 10 min at 4°C and the microsomal membranes were solubilized by incubation in the presence of 30 mM 1,2-diheptanoyl-sn-glycero-phosphocholine (DHPC, Avanti Polar Lipids) for 5 min at 4°C. Insoluble fraction was removed by centrifugation at 20,000 g for 10 min at 4°C. The resulting lysates were used to isolate total RNA using Direct-Zol RNA MiniPrep Plus (Zymo Research) or to affinity purify ribosomes with translated RNA using GFP-TRAP beads (magnetic agarose beads coated with Alpaca anti-GFP VHH antibody, Chromotek). Incubation with the GFP-TRAP beads proceeded for 3 h at 4°C with gentle end-over-end mixing in a tube rotator and was followed by three washes with high salt buffer (350 mM KCl, 10 mM MgCl_2_, 20 mM HEPES, 1% NP-40, 0.5 mM DTT, 100 μg/ml cycloheximide). Translated RNA was eluted from the beads by incubation in TRIZOL and isolated using Direct-Zol RNA MiniPrep Plus (Zymo Research).

RNA aliquots were stored at −70°C and only thawed once to minimize RNA degradation. RNA Quality Control analysis was performed using Agilent TapeStation 2200. Only samples with RNA Integrity Numbers >7 were used for subsequent analyses.

### RNA-Seq analyses

Total and translated RNA pools were used to prepare poly(A)-enriched cDNA libraries using standard protocol for BMDM and low input protocol for FLDM. Multiplex (8 samples/lane) paired-end (75 bp) sequencing was carried out using Illumina HiSeq 2500 at Yale Center for Genome Analyses. Raw sequencing data were trimmed for quality and aligned to the Mus musculus reference genome GRCm38 using HiSAT. The alignments were processed using ballgown, and per-gene counts were obtained. The raw counts were processed using DESeq2 and R.

Subsequent analyses used Transcripts Per kilobase Million (TPM) as a measure of mRNA abundance in total and translated poly(A) samples and Log2 fold change (Log2FC) to determine the effects of hypoxia within the related samples. Multiple comparisons with false-discovery rate (FDR) approach determined by the two-stage step-up method of Benjamini, Krieger and Yekutieli were applied to identify transcripts that significantly increased or decreased in response to hypoxia based on desired FDR Q=1%. Statistical analyses of the samples derived from BMDM and FLDM were performed separately. Only the protein coding transcripts showing statistically significant change of more than 1.2-fold (−0.32193<Log2FC<0.263034) in response to hypoxia were used for subsequent analyses. The resulting lists of transcripts differentially expressed or translated in response to hypoxia were displayed and analyzed using BioVenn web application (Hulsen et al., 2008) and Ingenuity Pathway Analyses (QIAGEN).

### Immunoblotting

BMDM and FLDM were differentiated and exposed to hypoxia as described above, rinsed with ice-cold PBS and lysed in 400 μl of lysis buffer containing 1% Igepal (Sigma) and complete protease inhibitors. Scraped lysates were precleared by centrifugation at 14,000 g for 10 min at 4°C. We separated proteins in 4-20% precast gradient gels (Bio-Rad) and transferred them onto nitrocellulose membrane. After 1 h blocking with the blocking buffer (Rockland), we probed the membranes sequentially with mouse anti-HuR (clone 3A2) and anti-actin primary antibodies (both from SantaCruz) and with goat anti-mouse AlexaFluor-680-conjugated secondary antibodies (ThermoFisher). We scanned the blots with LICOR Odyssey system and analyzed the intensity of the bands with ImageStudio Lite software version 5.2.

## Acknowledgments

Special thanks Drs. Chris Castaldi and Sameet Mehta at Yale Center for Genome Analyses who performed RNA-seq and provided bioinformatics support, respectively, and to Dr. Fatima Zahra Saddouk for assistance with the hypoxia protocol. We also thank faculty and staff at HHMI and in the Yale School of Medicine Office of Student Research (OSR), who provided essential administrative support. We acknowledge our funding sources, including the Howard Hughes Medical Institute (HHMI) Medical Research Fellowship (2017-2018, NSW), the Yale School of Medicine Marvin Moser Research Fellowship (2018, NSW), NIH/DHHS R01GM126412 (JRB) and the Connecticut Biomedical Research Fund (JRB).

